# Non-clonal emergence of colistin resistance associated with mutations in the BasRS two-component system in *Escherichia coli* bloodstream isolates

**DOI:** 10.1101/864983

**Authors:** Axel B. Janssen, Toby L. Bartholomew, Natalia P. Marciszewska, Marc J.M. Bonten, Rob J.L. Willems, Jose A. Bengoechea, Willem van Schaik

**Author notes:** Address correspondence to Willem van Schaik,.

## Abstract

Infections by multidrug-resistant Gram-negative bacteria are increasingly common, prompting the renewed interest in the use of colistin. Colistin specifically targets Gram-negative bacteria by interacting with the anionic lipid A moieties of lipopolysaccharides, leading to membrane destabilization and cell death. Here, we aimed to uncover the mechanisms of colistin resistance in nine colistin-resistant *Escherichia coli* strains and one *E. albertii* strain. These were the only colistin-resistant strains out of 1140 bloodstream *Escherichia* isolates collected in a tertiary hospital over a ten-year period (2006 - 2015). Core genome phylogenetic analysis showed that each patient was colonised by a unique strain, suggesting that colistin resistance was acquired independently in each strain. All colistin-resistant strains had lipid A that was modified with phosphoethanolamine. In addition, two *E. coli* strains had hepta-acylated lipid A species, containing an additional palmitate compared to the canonical hexa-acylated *E. coli* lipid A. One *E. coli* strain carried the mobile colistin resistance (mcr) gene *mcr-1.1* on an IncX4-type plasmid. Through construction of chromosomal transgene integration mutants, we experimentally determined that mutations in *basRS*, encoding a two-component signal transduction system, contributed to colistin resistance in four strains. We confirmed these observations by reversing the mutations in *basRS* to the sequences found in reference strains, resulting in loss of colistin resistance. While the *mcr*-genes have become a widely studied mechanism of colistin resistance in *E. coli*, sequence variation in *basRS* is another, potentially more prevalent but relatively underexplored, cause of colistin resistance in this important nosocomial pathogen.

**Importance:** Multidrug resistance among Gram-negative bacteria has led to the use of colistin as a last-resort drug. The cationic colistin kills Gram-negative bacteria through electrostatic interaction with the anionic lipid A moiety of lipopolysaccharides. Due to increased use in clinical and agricultural settings, colistin resistance has recently started to emerge. In this study, we used a combination of whole genome sequence analysis and experimental validation to characterise the mechanisms through which *E. coli* strains from bloodstream infections can develop colistin resistance. We found no evidence of direct transfer of colistin-resistant isolates between patients. The lipid A of all isolates was modified by the addition of phosphoethanolamine. In four isolates, colistin resistance was experimentally verified to be caused by mutations in the *basRS* genes, encoding a two-component regulatory system. Our data show that chromosomal mutations are an important cause of colistin resistance among clinical *E. coli* isolates.

## Introduction

*Escherichia coli* is a Gram-negative opportunistic pathogen that is a common cause of bloodstream, urinary tract, and enteric infections (1). The rising prevalence of antibiotic resistance in *E. coli*, in part due to the increasing global spread of the successful multidrug-resistant clade C lineage of ST131, may limit options for future treatments of infections (2, 3). Due to the emergence and spread of multidrug-resistant clones of *E. coli* and other Enterobacteriaceae, and the lack of new antibiotics targeting Gram-negative bacteria, colistin (polymyxin E) is increasingly used, despite its neuro- and nephrotoxic side effects, in the treatment of clinical infections with multidrug-resistant and carbapenem-resistant *E. coli* and other Enterobacteriaceae (4–6).

Colistin is a cationic, amphipathic molecule consisting of a non-ribosomal synthesized decapeptide and a lipid tail (7, 8). Colistin specifically targets Gram-negative bacteria by binding to the anionic phosphate groups of the lipid A moiety of lipopolysaccharides (LPS) through electrostatic interactions (7–9). Colistin destabilizes the outer membrane, but the subsequent disruption of the inner membrane ultimately leads to cell death (9, 10). Acquired colistin resistance has been reported in various Gram-negative bacteria that were isolated from clinical, veterinary, and environmental sources (11–13). The best-documented mechanism of colistin resistance involves the modification of lipid A with cationic groups to counteract the electrostatic interactions between colistin and lipid A (9). Lipid A modifications in Enterobacteriaceae may be mediated by the acquisition of mutations in chromosomally located genes or the acquisition of a mobile genetic element carrying one of the mobile colistin resistance (*mcr*)-genes, which encode phosphoethanolamine transferases that catalyse the addition of a cationic phosphoethanolamine group to lipid A (14–16).

Among Enterobacteriaceae, colistin resistance has been most intensively studied in *Salmonella* and *Klebsiella pneumoniae* in which mutations in the regulatory genes *mgrB, phoPQ* and *pmrAB* are important mechanisms leading to resistance (15, 17–19). In *E. coli* however, mutations in *mgrB* and *phoPQ* have not been reported to lead to colistin resistance. This may be caused by the increased rate of dephosphorylation of PmrA (BasR in *E. coli*) by PmrB (BasS in *E. coli*) in *E. coli* compared to other Enterobacteriaceae, which effectively negates the possible activating effects of mutations in *phoPQ* or *mgrB*, through PmrD, on the levels of phosphorylated BasR. This may explain why not all of the previously described mutations reported to lead to colistin resistance in *Salmonella* and *Klebsiella* confer resistance in *E. coli* (14, 20–22). In addition, *phoPQ* expression in *E. coli* is not only controlled by MgrB but also by the sRNA MicA, adding to the mechanisms controlling PhoPQ activation and making it less likely that the deletion or inactivation of *mgrB* can contribute to colistin resistance in *E. coli* (14, 23). This may explain why colistin resistance in clinical *E. coli* strains has only been linked to mutations in *basRS* (24–28), although experimental validation of the role of these mutations in colistin resistance is currently mostly lacking.

The PmrAB (BasRS) two-component system plays a crucial role in mediating the modification of LPS that lead to colistin resistance in Gram-negative bacteria (14, 17). Normally, this two-component system is activated by environmental stimuli, such as the presence of antimicrobial peptides or a low pH. Activation can increase virulence and survival through evasion of the host immune system by upregulating genes associated with modification of LPS, which is the predominant immunogenic molecule of Gram-negative bacteria (29, 30). In *E. coli*, the activation of BasRS leads to increased expression of various operons, including its own. This operon also includes *eptA*, which encodes a lipid A-specific phosphoethanolamine transferase (11, 14, 31).

Relatively little is known about colistin resistance mechanisms in *E. coli*, other than the acquisition of *mcr*-genes (32). Therefore, we studied a collection of colistin-resistant *E. coli* strains from bloodstream infections by a combination of whole genome sequencing and matrix-assisted laser desorption-ionisation time-of-flight (MALDI-TOF) analysis of their lipid A, to identify colistin resistance mechanisms in *E. coli*. The role of mutations in *basRS* was investigated through the construction of chromosomal integration mutants of different *basRS* alleles.

## Materials and methods

### Ethical statement

Approval to obtain data from patient records was granted by the Medical Ethics Review Committee of the University Medical Center Utrecht, in Utrecht, The Netherlands (project numbers 16/641 and 18/472).

Colistin-resistant *E. coli* strains were isolated as part of routine diagnostic procedures. This aspect of the study did not require consent or ethical approval by an institutional review board.

### Bacterial strains, growth conditions, and chemicals

Colistin-resistant *E. coli* strains from bloodstream infections were obtained retrospectively from the strain collection of the clinical microbiology laboratory of the University Medical Center Utrecht in Utrecht, The Netherlands. In initial routine diagnostic procedures, blood cultures were plated on TSA plates with 5% sheep blood. Strains collected up to 2011 were identified and their antibiogram was determined using the BD Phoenix automated identification and susceptibility testing system (Becton Dickinson, Vianen, The Netherlands). From 2011 onwards, species determination was performed by MALDI-TOF on a Bruker microflex system (Leiderdorp, The Netherlands). *E. coli* strain BW25113 and the BW25113-derived Δ*basRS* strain BW27848 from the Keio collection were obtained from the Coli Genetic Stock Center (33, 34). Strains were grown in Lysogeny Broth (LB; Oxoid, Landsmeer, The Netherlands) at 37°C with agitation at 300 rpm unless otherwise noted, with exception of strains containing pGRG36, which were grown at 30°C (35). When appropriate, kanamycin (50 mg/L; Sigma-Aldrich, Zwijndrecht, The Netherlands), and ampicillin (100 mg/L; Sigma-Aldrich) were used. Colistin sulphate was obtained from Duchefa Biochemie (Haarlem, The Netherlands). L-(+)-arabinose was obtained from Sigma-Aldrich. Plasmids were purified using the GeneJET Plasmid Miniprep kit (Thermo Fisher Scientific, Landsmeer, The Netherlands). PCR products were purified from gel using GeneJET Gel Extraction and DNA Cleanup Micro Kit (Thermo Fisher Scientific).

### Determination of minimal inhibitory concentration

Minimal inhibitory concentrations (MICs) to colistin were determined as previously described (36) in line with the recommendations of a joint working group of the Clinical & Laboratory Standards Institute and the European Committee on Antimicrobial Susceptibility Testing (EUCAST) (http://www.eucast.org/fileadmin/src/media/PDFs/EUCAST_files/General_documents/Recommendations_for_MIC_determination_of_colistin_March_2016.pdf), using BBL™ Mueller Hinton II (cation-adjusted) broth (MHCAB; Becton Dickinson), untreated Nunc 96-wells round bottom polystyrene plates (Thermo Scientific), and Breathe-Easy sealing membranes (Sigma-Aldrich). The breakpoint value of an MIC > 2 µg/ml for colistin resistance in *E. coli* was obtained from EUCAST (http://www.eucast.org/clinical_breakpoints/).

### Genomic DNA isolation and whole-genome sequencing

Genomic DNA was isolated using the Wizard Genomic DNA purification kit (Promega, Leiden, The Netherlands) according to the manufacturer’s instructions. DNA concentrations of the genomic DNA preparations were measured with the Qubit dsDNA Broad Range Assay kit and the Qubit 2.0 fluorometer (Life Technologies, Bleiswijk, The Netherlands) and were all higher than 20 ng/µl.

Sequence libraries for Illumina sequencing were prepared using the Nextera XT kit (Illumina, San Diego, CA) according to the manufacturer’s instructions with 1 ng genomic DNA as input. Libraries were sequenced on an Illumina MiSeq system with a 500-cycle (2 × 250 bp) MiSeq reagent kit v2.

For strain E3090, we performed long-read sequencing using the MinION platform (Oxford Nanopore Technologies) to fully resolve the *mcr-1.1* plasmid. MinION library preparation for barcoded 2D long-read sequencing was performed using the SQK-LSK208 kit (Oxford Nanopore Technologies, Oxford, England, United Kingdom), according to the manufacturer’s instructions, with G-tube (Covaris, Woburn, Massachusetts, United States of America) shearing of 1 µg chromosomal DNA for 2 x 120 seconds at 1500 *g*. Sequencing was performed on the MinION sequencer (Oxford Nanopore Technologies) using 2D barcoded sequencing through a SpotON Flow Cell Mk I (R9.4; Oxford Nanopore Technologies).

### Genome assembly, MLST typing, and identification of antibiotic resistance genes

The quality of Illumina sequence data was assessed using FastQC v0.11.5 (https://github.com/s-andrews/FastQC). Raw Illumina sequencing reads were trimmed for quality using nesoni v0.115 (https://github.com/Victorian-Bioinformatics-Consortium/nesoni) using standard settings with the exception of a minimum read length of 100 nucleotides. *De novo* genome assembly of the trimmed Illumina short-read data was performed using SPAdes v3.6.2 with the following settings: kmers used: 21, 33, 55, 77, 99, or 127, “careful” option turned on and cut-offs for final assemblies: minimum contig/scaffold size = 500 bp, minimum contig/scaffold average Nt coverage = 10-fold (37).

MinION sequence read data in FastQ format was extracted from Metrichor base-called raw FAST5 read-files using Poretools (38). A hybrid assembly for strain E3090 was generated with trimmed Illumina short-read data and Oxford Nanopore Technologies MinION long-read data by using SPAdes v3.6.2 with the same settings as the Illumina short-read assemblies, and specifying the long-read data with the --nanopore flag.

Gene prediction and annotation was performed using Prokka (39), using standard settings. MLST typing was performed using the mlst package v2.10 (https://github.com/tseemann/mlst), using standard settings. Assembled contigs were assessed for antibiotic resistance genes using ResFinder 3.2 (40), using standard settings.

### Core genome phylogenetic analysis and determination of mutations in candidate colistin resistance determinants

Genome assemblies generated in this study with Illumina data were aligned with 178 complete *E. coli* genomes and 32 *E. albertii* genomes that were available from NCBI databases on 24 June 2016 (Supplemental Table S1) using ParSNP v1.2 (41). MEGA6 was used to midpoint root and visualize the phylogenetic tree (42). We identified whether non-synonymous mutations were present in *basRS* by pairwise comparison of the gene sequences of colistin-resistant isolates to their closest matching publicly available genome from the phylogenetic tree using BLAST (43). Mutations that were identified in the genome sequences were confirmed through PCR (oligonucleotide primer sequences are provided in Supplemental Table S2) and subsequent Sanger sequencing of the PCR product by Macrogen (Amsterdam, The Netherlands).

### Isolation and analysis of lipid A

Isolation of lipid A molecules and subsequent analysis by negative-ion MALDI-TOF mass spectrometry was performed as previously described (19, 44, 45). Briefly, *Escherichia* strains were grown in LB (Oxoid) and the lipid A was purified from stationary cultures using the ammonium hydroxide/isobutyric acid method described earlier (46). Mass spectrometry analyses were performed on a Bruker autoflex™ speed TOF/TOF mass spectrometer in negative reflective mode with delayed extraction using as matrix an equal volume of dihydroxybenzoic acid matrix (Sigma-Aldrich) dissolved in (1:2) acetonitrile-0.1% trifluoroacetic acid. The ion-accelerating voltage was set at 20 kV. Each spectrum was an average of 300 shots. A peptide calibration standard (Bruker) was used to calibrate the MALDI-TOF. Further calibration for lipid A analysis was performed externally using lipid A extracted from *E. coli* strain MG1655 grown in LB medium at 37°C.

### Construction of chromosomal *basRS* transgene insertions

Chromosomal transgene insertions of *basRS* were constructed in BW27848 by utilizing the Tn7 transposon system on the pGRG36 plasmid (35). The strategy to generate the constructs is illustrated in Supplemental Figure S1. The promoter of the *eptA*-*basRS* operon was fused to the *basRS* coding sequence by separate PCRs for the promotor region and the *basRS* amplicon, with high fidelity Phusion Green Hot Start II DNA Polymerase (Thermo Fisher Scientific) using strain-specific primers (Supplemental Table S2; oligonucleotides were obtained from Integrated DNA Technologies, Leuven, Belgium). The promoter and the *basRS* amplicon were subsequently fused by overlap PCR. Fused PCR products were cloned into pCR-Blunt II-TOPO using the Zero Blunt TOPO PCR Cloning kit (Thermo Fisher Scientific), and subsequently subcloned into pGRG36 (35). Electrocompetent BW25113 and BW27848 *E. coli* cells were prepared as described previously (47) and transformed using the following settings: voltage 1800V, capacitance 25 µF, resistance 200Ω, with a 0.2 cm cuvette using the Gene Pulser Xcell electroporation system (Bio-Rad Laboratories, Veenendaal, The Netherlands). Transformants were grown at 30°C. After confirming integration of the Tn7 transposon at the *att*Tn7 site by PCR (primers listed in Supplemental Table S2) and Sanger sequencing (Macrogen), the pGRG36 plasmid was cleared by culturing at 37°C.

Inverse PCR site-directed mutagenesis was performed on amplicons cloned in pCR-Blunt II-TOPO to reverse the mutations that were identified in colistin-resistant strains to the sequences of *basR* or *basS* in the closest matching publicly available genome (48). After gel purification of the amplified fragments, (hemi)methylated fragments were digested using DpnI (New England Biolabs (NEB), Ipswich, Massachusetts, United States of America). Subsequently the vector was recircularized using the Rapid DNA Ligation kit (Thermo Fisher Scientific) after phosphorylation using T4 Polynucleotide kinase (NEB). The constructs were then transformed into chemically competent DH5α *E. coli* cells (Invitrogen, Landsmeer, The Netherlands). Mutated sequences were subsequently subcloned to pGRG36 as described above.

## Data availability

Sequence data has been deposited in the European Nucleotide Archive (accession number PRJEB27030).

## Results

### Low prevalence of colistin resistance in invasive *Escherichia* bloodstream isolates

A total of 1140 bloodstream isolates (collected from January 2006 to December 2015) for which species identification had been performed and automated antibiotic susceptibility profiles had been determined, were available for this study. Twelve isolates were deemed resistant to colistin through routine diagnostic procedures. Two of those isolates were isolated from the same patient, on the same day, and were thus considered duplicates, and only one of these was included in this study. In ten of the eleven remaining isolates, colistin resistance, defined as an MIC > 2 µg/ml colistin, was confirmed through broth microdilution (Table 1). Strain A783 was a false positive for colistin resistance during automated susceptibility testing in routine diagnostic procedures, and was excluded from subsequent analyses, leaving ten isolates for further investigation.

**Table 1:** Colistin-resistant *Escherichia* strains isolated from bloodstream infections. Overview of colistin-resistant bloodstream isolates, including the MIC of colistin, MLST type determined through whole genome sequencing, date of isolation, and information on the use of colistin three months before the isolation of the colistin-resistant isolate, and if applicable, route of administration. The MIC values represent the median of three independent replicate experiments performed in triplicate.

| Strain | Colistin MIC (µg/ml) | MLST | Date of isolation | History of colistin use |
| --- | --- | --- | --- | --- |
| **I1121** | 16 | 131 | 22 April 2015 | Yes; inhalation and oral |
| **H2129** | 8 | 131 | 22 July 2014 | No |
| **G821** | 16 | 131 | 19 March 2013 | No |
| **F2745** | 4 | 73 | 2 November 2012 | No |
| **E3090** | 8 | 10 | 12 November 2011 | No |
| **E2372** | 4 | 59 | 25 August 2011 | No |
| **E650** | 8 | 162 | 11 March 2011 | No |
| **D2373** | 8 | 6901 | 20 October 2010 | Yes; oral |
| **A2361** | 8 | 5268 | 3 November 2007 | No |
| **Z821** | 4 | 167 | 2 April 2006 | Yes; oral |

The estimated prevalence of colistin resistance in *E. coli* strains causing bloodstream infections isolated from January 2006 to December 2015 was thus determined to be 0.88%. Three patients had received colistin in the three months before isolation of the colistin-resistant strain (Table 1). Two of these patients received colistin to treat infections, but all three patients were also administered colistin as part of selective digestive or oropharyngeal decontamination (SDD/SOD), a prophylactic antibiotic treatment widely used in Dutch intensive care units (49). The ten colistin-resistant strains were analyzed further in this study to determine their relatedness and mechanism through which they had developed colistin resistance.

### Colistin resistance was independently acquired by each individual bloodstream *E. coli* isolate

To assess the phylogenetic relationships between the colistin-resistant strains, a phylogenetic tree was generated based on the genome assemblies of the colistin-resistant strains and 210 publicly available complete genome sequences (Supplemental Table S1). Based on a core genome alignment of 874 kbp, we did not observe direct transmission of colistin-resistant strains between patients (Figure 1A). Three colistin-resistant strains (strains I1121, H2129, and G821) belonged to the globally disseminated ST131 clone, and all three were dispersed throughout the multidrug-resistant clade C of ST131(Figure 1A, 1B) (3, 50). This indicates that the ST131 strains in this study have independently acquired colistin resistance. Strain A2361 clustered among *E. albertii* (Figure 1A), although it had been typed as *E. coli* in routine diagnostic procedures.

**Figure 1:**
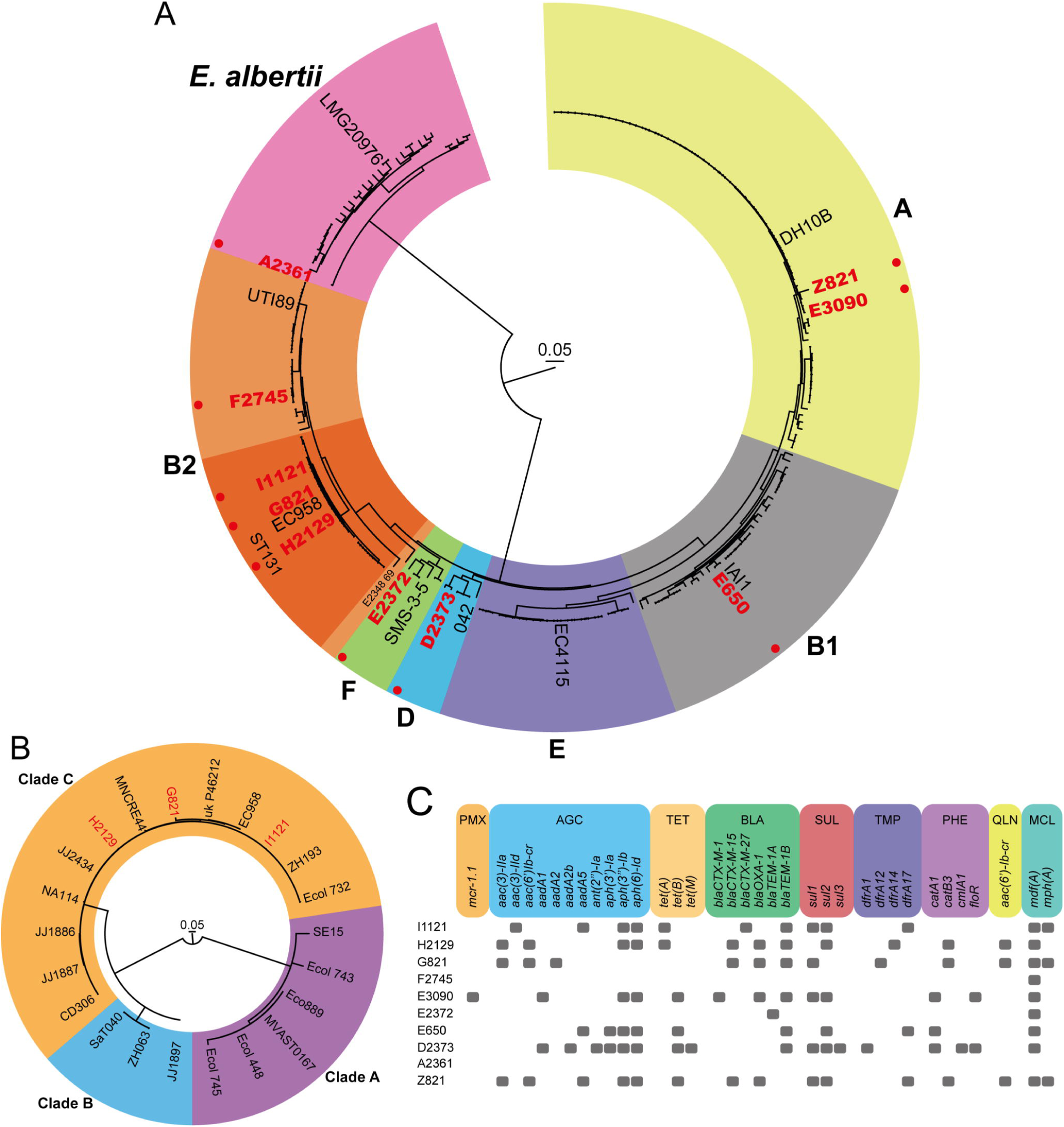
Colistin-resistant strains are not clonally related, and carry diverse acquired antibiotic resistance genes. **A)** The phylogenetic tree represents the core genome alignment (874 kbp) of the colistin-resistant strains and 210 publicly available *E. coli* and *E. albertii* genome sequences. One representative reference strain per *E. coli* phylogroup is indicated (64). For *E. albertii*, the LMG20976 type-strain is indicated (65). The different phylogroups of *E. coli* are indicated with coloured backgrounds. Phylogroup A is coloured yellow; B1, grey; B2, orange; D, blue; E, purple; F, green. The ST131 lineage of *E. coli* in phylogroup B2 is indicated by a dark orange background. The *E. albertii* branch is indicated by a pink background. The colistin-resistant strains characterized in this study are depicted in red, and highlighted by a red filled circle. **B)** The phylogenetic tree represents the core genome alignment (3.55 Mbp) of the three colistin-resistant ST131 strains and 19 publicly available ST131 *E. coli* strains genome sequences. The colistin-resistant strains characterized in this study are depicted in red. Clades A, B, and C of ST131 are indicated by purple, blue, and orange backgrounds respectively. **C)** Antibiotic resistance genes in the genome sequences were detected by ResFinder 3.2 (40). Classes of antibiotic resistance genes are abbreviated as follow: PMX, polymyxin resistance; AGC, aminoglycoside resistance; TET, tetracycline resistance; BLA, beta-lactam resistance; SUL, sulfonamide resistance; TMP, trimethoprim resistance; PHE, phenicol resistance; QLN, quinolone resistance; MCL, macrolide.

By screening for acquired antibiotic resistance genes through ResFinder 3.2, we found that only strain E3090 carried the *mcr* gene *mcr-1.1* (0.086% of all bloodstream isolates; Figure 1C). After long-read sequencing and hybrid assembly, the *mcr-1.1*-gene in this strain appeared to be located as the sole antibiotic resistance gene on a 32.7 kbp IncX4-type plasmid. This *mcr-1.1* carrying IncX4-type plasmid from E3090 shares 99% identity to the previously reported *mcr-1.1* carrying IncX4-type plasmid pMCR-1_Msc (GenBank accession MK172815.1) harboured by *E. coli* isolated from patients in Russia (51), confirming the global dissemination of this plasmid (52). In all strains studied here, a variety of acquired resistance genes was observed (Figure 1C), reflecting the non-clonal nature of the colistin-resistant strains. The three colistin-resistant ST131 strains possessed different repertoires of acquired resistance genes, further excluding recent transmission between patients of the ST131 strains studied here. Strain F2745 and E2372 carried only one, and two resistance genes respectively, while the *E. albertii* strain A2361 did not possess any acquired resistance genes.

### *Escherichia* isolates exclusively acquire colistin resistance by modification of phosphate groups of lipid A

To determine which modifications to lipid A are affecting colistin resistance in *E. coli* we extracted lipid A from the clinical strains and the colistin-susceptible control *E. coli* strain MG1655, and subjected them to MALDI-TOF mass spectrometry. The lipid A produced by all *E. coli* strains showed lipid A species with a mass-to-charge ratio (*m/z)* of 1797 (Figure 2A), corresponding to the canonical unmodified *E. coli* hexa-acylated lipid A (Figure 2B). Colistin-resistant strains showed additional lipid A species at *m/z* 1921, consistent with the addition of phosphoethanolamine (*m/z* 124) to the hexa-acylated species. Additional species were detected in the lipid A produced by strains E650 and Z821. Species *m/z* 2036 indicated the addition of palmitate (*m/z* 239) to the hexa-acylated species *m/z* 1797, whereas species *m/z* 2160 was consistent with the addition of palmitate to the hexa-acylated lipid A species containing phosphoethanolamine (*m/z* 1910).

**Figure 2:**
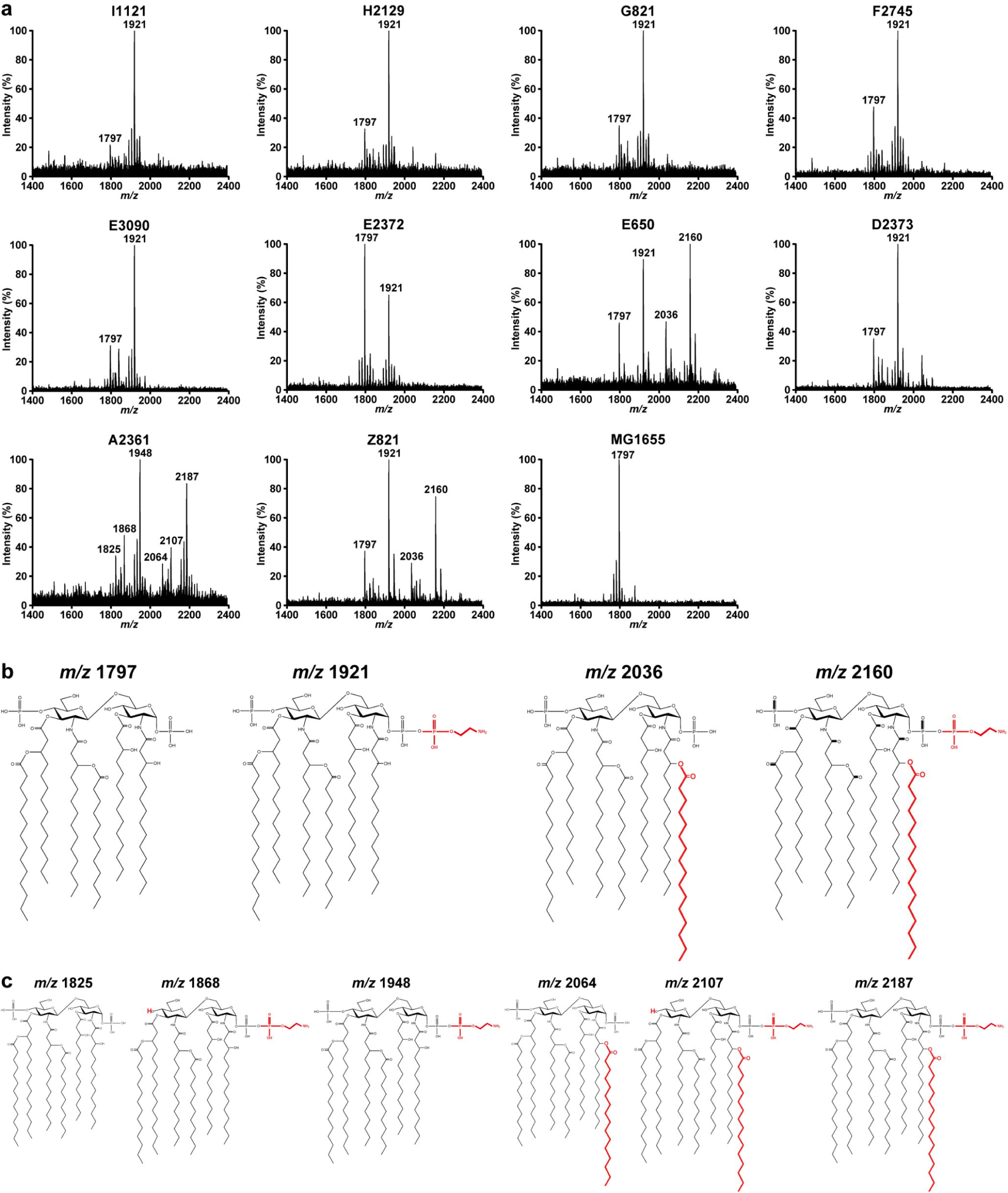
MALDI-TOF spectra of lipid A from colistin-resistant nosocomial *Escherichia* strains. Negative ion MALDI-TOF mass spectrometry spectra of lipid A purified from **A)** colistin-resistant strains and colistin-susceptible MG1655. Data represent the mass to charge (*m/z*) ratios of each lipid A species detected and are representative of three extractions. **B)** Proposed lipid A structures of the species produced by *E. coli* strains. **C)** Proposed lipid A structures of *E. albertii* strain A2361. Modifications to unmodified lipid A are depicted in red.

The *E. alberti* strain A2361 produced lipid A distinct from *E. coli.* Species *m/z* 1825 is likely to represent a hexa-acylated species corresponding to two glucosamines, two phosphates, four 3-OH-C_14_, and two C_14_ (Figure 2C) Species *m/z* 1948 is consistent with the addition of phosphoethanolamine to the hexa-acylated species, with a further addition of palmitate to produce lipid A species *m/z* 2187. Species *m/z* 1868 and *m/z* 2107 could correspond to the loss of the second phosphate group, compared to *m/z* 1948, and *m/z* 2187.

### Identification of mutations in *basRS* as candidate mutations involved in colistin resistance

Because chromosomal mutations in *basRS*, but not in other regulatory systems, have previously been suggested to cause colistin resistance in *E. coli* (24–28), we next aimed to establish the contribution of the *basRS* alleles in the colistin-resistant phenotype of these bloodstream isolates. Due to the multidrug-resistant nature of the clinical isolates (Figure 1C), we were unable to generate targeted mutations in these strains. Therefore, we made chromosomal transgene insertion mutants of the different *basRS* alleles in the *att*Tn7 site in the BW25113-derived Δ*basRS* strain BW27848 using the Tn7 transposon system. By making chromosomal transgenes insertions, rather than using an *in trans* complementation method, we excluded copy number effects by plasmids, and the need to use antibiotics to select for the presence of a plasmid used for *in trans* complementation. Since BW27848 still possesses the gene encoding for the phosphoethanolamine transferase EptA, we constructed sequences that consisted of the fused sequences of the promotor region of the *eptA*-*basR*-*basS* operon and the *basRS* coding sequences in order to prevent *eptA* gene dosage-dependent effects. We were unable to generate the construct for strain E650, presumably due to the toxicity of the insert.

The colistin MIC determination of the generated *basRS* chromosomal transgene insertion mutants from strains I1121, H2129, G821, and Z821 had higher colistin MIC values than the BW27848::Tn7-empty strain, with observed MIC values ≥16-fold higher than that of the BW27848::Tn7-empty strain (Table 2). As expected, the *basRS* allele of the *mcr-1.1* positive strain E3090 did not lead to colistin resistance. We were unable to show the contribution of *basRS* to colistin resistance in the additional four colistin-resistant strains (F2745, E2372, D2373, A2361) that lacked *mcr*-*1.1*.

**Table 2.** MICs of colistin of strains generated in this study. *E. coli* strain BW27848 is the Δ*basRS* mutant of BW25113 (34). The *basRS* alleles of colistin-resistant strains from this study were inserted into the *att*Tn7 site of BW27848. The addition of “m” to a strain name indicates that the construct has been modified through inverse PCR site-directed mutagenesis to reverse the mutation associated with colistin resistance. The values represent the median of three independent replicate experiments performed in triplicate.

| Strain | Colistin MIC<br>($\mu\text{g/ml}$) |
| --- | --- |
| BW25113 | 0.25 |
| BW27848 | 0.125 |
| BW25113::Tn7 Empty | 0.25 |
| BW27848::Tn7 Empty | 0.125 |
| BW27848::Tn7 BW25113 | 0.125 |
| BW27848::Tn7 I1121 | 2 |
| BW27848::Tn7 I1121m | 0.25 |
| BW27848::Tn7 H2129 | 4 |
| BW27848::Tn7 H2129m | 0.25 |
| BW27848::Tn7 G821 | 4 |
| BW27848::Tn7 G821m | 0.25 |
| BW27848::Tn7 F2745 | 0.25 |
| BW27848::Tn7 E3090 | 0.125 |
| BW27848::Tn7 E2372 | 0.25 |
| BW27848::Tn7 D2373 | 0.5 |
| BW27848::Tn7 A2361 | 0.125 |
| BW27848::Tn7 Z821 | 2 |
| BW27848::Tn7 Z821m | 0.125 |

### Mutations in the *basRS* genes contribute to colistin resistance in *E. coli*

By construction of the chromosomal transgene insertion mutants, we identified the ability of the *basRS* sequences of four strains (I1121, H212, G821, and Z821) to cause colistin resistance in BW27848. To identify the mutations in the *basRS* alleles of these strains that contribute to resistance, we compared the *basRS* encoding sequences of those strains causing resistance to the phylogenetically most closely related publicly available *E. coli* genome sequences used in the construction of Figure 1A. None of these reference strains were reported to be colistin-resistant, or carried any of the *mcr*-genes. This comparison revealed four distinct mutations: a L10R substitution in BasS in I1121, a G53S substitution in BasR in H2192, the duplication of the HAMP-domain in BasS in G821, and a A159P substitution in BasS in Z821 (Supplemental Figure S2). As expected, in the *mcr-1.1* positive strain E3090 no mutations in *basRS* were identified.

We hypothesised that the observed mutations were impacting the normal functioning of the BasRS two-component system. To assess whether the mutations in *basRS* identified by comparing the *basRS* sequences of the clinical strains I1121, H2129, G821, and Z821, and their closest match in the set of 178 publicly available *E. coli* genome sequences (Supplemental Figure S2) were causal to the development of colistin resistance, the identified mutations were reversed through site-directed inverse PCR mutagenesis to match the *basRS* alleles of the publicly available genome sequences. The MIC values of these mutants returned to levels similar to that of the colistin-susceptible BW27848::Tn7-empty strain (Table 2). These experiments support the involvement of *basRS* sequence variation in colistin resistance in *E. coli*.

## Discussion

In the present study, we set out to characterise the the mechanisms through which *E. coli* bloodstream isolates can develop colistin resistance through a combination of whole genome sequence analysis and experimental validation. We did not find evidence for transfer of colistin-resistant strains between patients, suggesting that colistin resistance has been acquired independently in all cases. In seven patients colistin-resistant strains were isolated without the patients being previously exposed to the drug. All colistin-resistant strains had LPS that was modified by the addition of phosphoethanolamine to the lipid A moiety of LPS. Resistance in one of the bloodstream isolates could be explained by the acquisition of *mcr-1.1*. In four other strains, we identified mutations in *basRS* that contribute to colistin resistance. Although colistin-susceptible strains that were isogenic to the resistant strains were not available, we were able to pinpoint the mutations in *basRS* leading to resistance in these strains by matching the genomic sequences of our nosocomial isolates with publicly available genomes, none of which were reported to be colistin-resistant, and subsequent construction of chromosomally integrated *basRS* transgene alleles in the Δ*basRS* strain BW27848. The mechanisms of colistin resistance in the remaining five strains remain to be characterized.

Some of the mutations we experimentally link to colistin resistance in this study, have previously been associated with colistin resistance or the functioning of the BasRS two-component system. In this study, we demonstrated that the amino acid change L10R in BasS (strain I1121) also confers colistin resistance. An amino acid substitution in the same position of BasS (L10P) was previously experimentally proven to cause colistin resistance in *E. coli* (26). The glycine in position 53 of BasR has previously been reported to be altered in colistin-resistant Enterobacteriaceae (53, 54) including in *E. coli* (55). The G53S change specifically, as in isolate H2192, has been experimentally proven to contribute to colistin resistance in *Klebsiella* (previously *Enterobacter*) *aerogenes* (56, 57) and *Salmonella enterica* subsp. *enterica* serovar Typhimurium (58) and we extend those findings to *E. coli* here. The previously unidentified duplication of 162 nucleotides in *basS* (strain G821) leads to the introduction of a second HAMP domain in BasS and confers colistin resistance in the BW27848 background. The HAMP domain is widespread in bacteria and is commonly involved in signal transduction as part of two-component systems (59). We hypothesise that the addition of an extra HAMP domain in BasS may change signal transduction in the protein, leading to the constitutive activation of the histidine kinase domain of BasS, increased phosphorylation of BasR and upregulated expression of *eptA*, ultimately resulting in the addition of phosphoethanolamine to lipid A. Finally, we demonstrate that the A159P substitution in BasS (observed in strain Z821) contributes to colistin resistance. A mutation leading to a A159V substitution was found in an *in vitro* evolution study in which *E. coli* was evolved towards colistin resistance (60), and in clinical colistin-resistant *E. coli* isolates (61), but experimental confirmation of the role of alterations in A159 in colistin resistance in *E. coli* was so far lacking. Our data suggest that the *basRS* alleles of three *E coli* strains (F2745, E2372, and D2373), and the *E. albertii* strain A2361, do not confer resistance in the BW25113 *E. coli* background. Because *E. albertii* is phylogenetically distinct from *E. coli*, its *basRS* allele may not function optimally in an *E. coli* background, explaining the inability of the transgene insertion complementation in the *basRS* deletion of BW25113 *E. coli* strain to cause colistin resistance (62). We are unable to explain the colistin resistance mechanisms of the clinical isolates F2745, E2372, and D2373. It is likely that these strains have become resistant to colistin through other mutations that finally lead to the modification of lipid A by phosphoethanolamine.

The observed modification of lipid A with phosphoethanolamine in all isolates underlines the crucial role of phosphoethanolamine transferases in the ability of *Escherichia* to become resistant to polymyxins (14). The lipid A of three of the colistin-resistant strains was also modified with palmitate, but the contribution of lipid A palmitoylation to colistin resistance in clinical *E. coli* strains is currently unknown. We did not observe modifications of lipid A by 4-amino-4-deoxy-L-arabinose in the colistin-resistant isolates. While this modification was shown to contribute to polymyxin B resistance under low Mg^2+^ conditions in a laboratory isolate of *E. coli* (20), it may be rare in clinical *E. coli* isolates. Indeed, Sato *et al*. also exclusively found phosphoethanolamine-modified lipid A in colistin-resistant clinical *E. coli* isolates (24). The reliance of *Escherichia* on the modification of lipid A by phosphoethanolamine to acquire colistin resistance, suggests that the inhibition of this class of enzymes by blocking the conserved catalytic site (31) could be a target for future drug development and opens the possibility of combination therapy with colistin and an inhibitor of phosphoethanolamine transferase (63). With the increasing clinical issues posed by infections with multidrug-resistant Gram-negative bacteria, there is an urgent need to better understand resistance mechanisms to last-resort antibiotics like colistin. While the discovery of the *mcr-*genes have generated considerable interest in transferable colistin resistance genes, our data suggest that chromosomal mutations remain an important cause of colistin resistance among clinical isolates in the genus *Escherichia*.

## Acknowledgements

We thank Eline A.M. Majoor for technical support and L. Marije Hofstra and Lidewij W. Rümke for their review of patient records. We also thank the Utrecht Sequence Facility and Ivo Renkens for their expertise in MinION Nanopore sequencing.

W.v.S. was funded through an NWO-Vidi grant (grant 917.13.357), and a Royal Society Wolfson Research Merit Award. Work in J.A.B. laboratory was supported by Biotechnology and Biological Sciences Research Council (BBSRC, BB/P020194/1) and Queen’s University Belfast start-up. T.L.B. is the recipient of a PhD fellowship funded by the Department for Employment and Learning (Northern Ireland, UK). The funders had no role in study design, data collection and interpretation, or the decision to submit the work for publication.

A.B.J. conceived and designed experiments, performed experiments, analysed data, and wrote the manuscript. T.B.L. performed experiments, and analysed data. N.P.M. performed experiments, and analysed data. M.J.M.B. wrote the manuscript. R.J.L.W. wrote the manuscript. J.A.B. analysed data, and wrote the manuscript. W.v.S. conceived and designed experiments, wrote the manuscript, and supervised the study. All authors reviewed and approved the final version of the manuscript.

## Competing interests

The authors declare no conflicts of interest.

