## Supplemental Figure S2 for "Non-clonal emergence of colistin resistance associated with mutations in the BasRS two-component system in *Escherichia coli* bloodstream isolates"

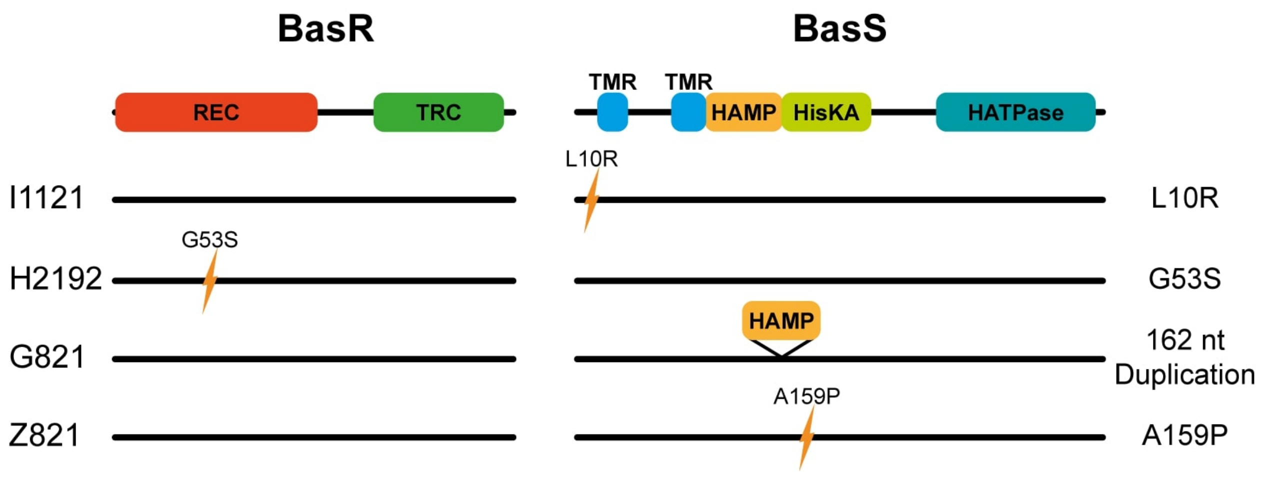


**Supplemental Figure S2: Conservation and prediction of functional effects of mutations in *basRS*.**

Comparison of the *basRS* sequences of colistin-resistant strains and publicly available genome sequences led to the identification of mutations in *basRS* that could have a role in colistin resistance. Domains of BasR and BasS were predicted using SMART (S1). The domains are REC: CheY-homologous receiver domain, TRC: Transcriptional regulatory protein, C terminal (Trans_reg_c), HAMP: Histidine kinases, Adenylyl cyclases, Methyl binding proteins, Phosphatases domain, HisKA: His Kinase A (phosphoacceptor) domain, HATPase: Histidine kinase-like ATPases (HATPase_c). The two transmembrane regions (TMR) in BasS are highlighted in blue.
