## Supplemental Figure S1 for "Non-clonal emergence of colistin resistance associated with mutations in the BasRS two-component system in *Escherichia coli* bloodstream isolates"

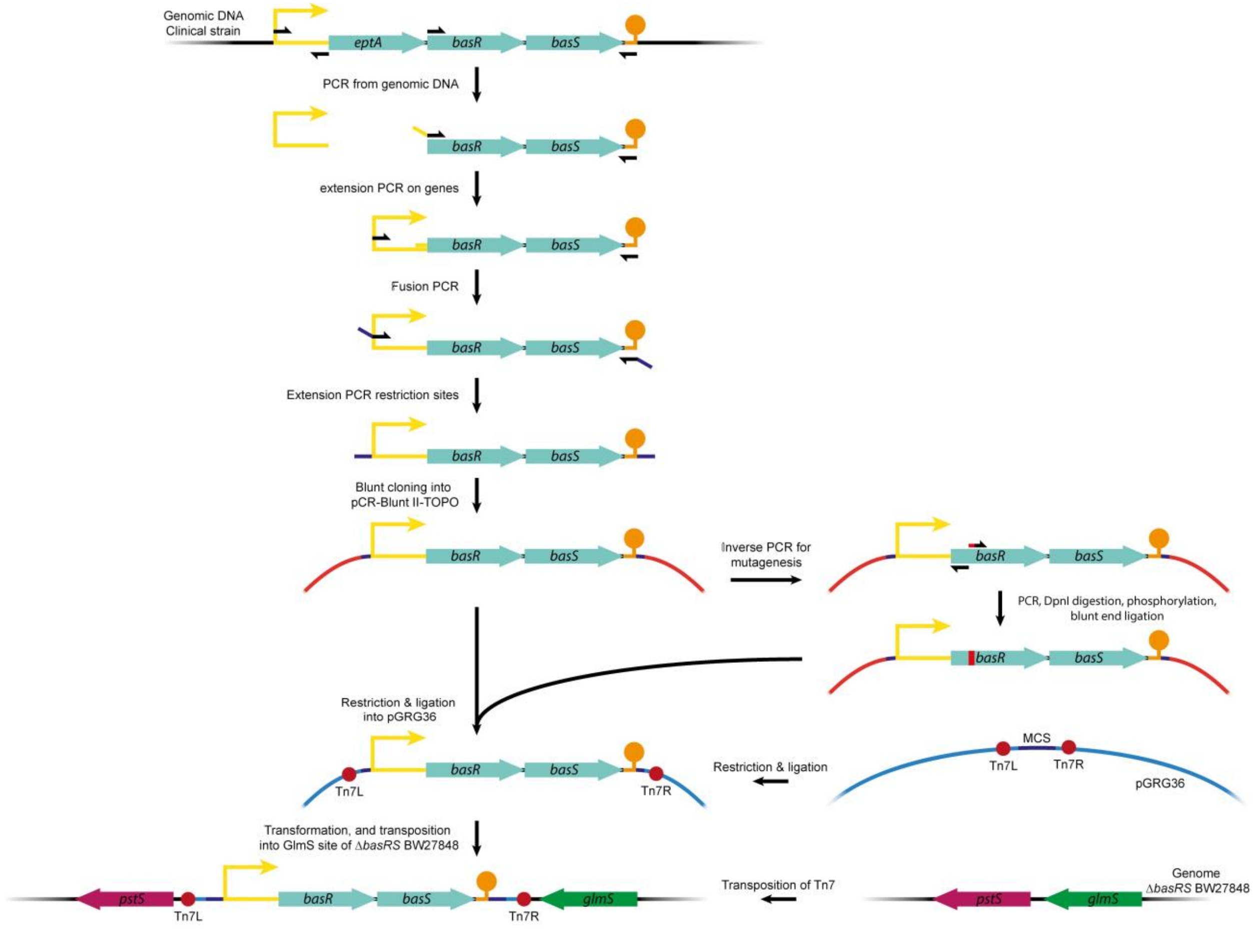


**Supplemental Figure S1: Strategy of the construction of chromosomal *basRS* transgene insertions.**

Chromosomal *basRS* transgene insertion mutants were inserted in the *att*Tn7 site in *E. coli* BW27848 (a Δ*basRS* derivative of BW25113). Genes encoded in the same operon are colored in the same color. Promoter, and terminator sequences are indicated, and colored individually. The position of the primers used in the construction of the constructs are indicated with half-arrows.
