## Supplemental Table S1 for "Non-clonal emergence of colistin resistance associated with mutations in the BasRS two-component system in *Escherichia coli* bloodstream isolates"

| **Strain** | **MLST** | **GenBank assembly accession number** | **Organism** | **Closest match** |
| --- | --- | --- | --- | --- |
| 42 | 414 | GCA_000027125.1 | *Escherichia coli* |  |
| 536 | 127 | GCA_000013305.1 | *Escherichia coli* |  |
| 789 | 88 | GCA_000819645.1 | *Escherichia coli* |  |
| 1303 | 10 | GCA_000829985.1 | *Escherichia coli* |  |
| 6409 | 10 | GCA_000814145.2 | *Escherichia coli* | E3090 |
| 11128 | 16 | GCA_000010765.1 | *Escherichia coli* |  |
| 11368 | 21 | GCA_000091005.1 | *Escherichia coli* |  |
| 12009 | 17 | GCA_000010745.1 | *Escherichia coli* |  |
| 55989 | 3762 | GCA_000026245.1 | *Escherichia coli* |  |
| 180‑PT54 | 5638 | GCA_001650275.1 | *Escherichia coli* |  |
| 2009C‑3133 | 117 | GCA_001420955.1 | *Escherichia coli* |  |
| 2009EL‑2050 | 678 | GCA_000299255.1 | *Escherichia coli* |  |
| 2009EL‑2071 | 678 | GCA_000299475.1 | *Escherichia coli* |  |
| 2011C‑3493 | 678 | GCA_000299455.1 | *Escherichia coli* |  |
| 2011C‑3911 | 1727 | GCA_001644725.1 | *Escherichia coli* |  |
| 2012C‑4227 | 119 | GCA_001420935.1 | *Escherichia coli* |  |
| 2013C‑4465 | 335 | GCA_001644745.1 | *Escherichia coli* |  |
| 28RC1 | 11 | GCA_001612475.1 | *Escherichia coli* |  |
| 644‑PT8 | N.D. | GCA_001650295.1 | *Escherichia coli* |  |
| 94‑3024 | 672 | GCA_000801185.2 | *Escherichia coli* |  |
| ABU 83972 | 73 | GCA_000148365.1 | *Escherichia coli* |  |
| ACN001 | 23 | GCA_001051135.1 | *Escherichia coli* |  |
| ACN002 | 23 | GCA_001515725.1 | *Escherichia coli* |  |
| APEC IMT5155 | N.D. | GCA_000813165.1 | *Escherichia coli* |  |
| APEC O1 | 95 | GCA_000014845.1 | *Escherichia coli* |  |
| APEC O78 | 23 | GCA_000332755.1 | *Escherichia coli* |  |
| ATCC 25922 | 73 | GCA_000743255.1 | *Escherichia coli* | F2745 |
| ATCC 8739 | 3021 | GCA_000019385.1 | *Escherichia coli* |  |
| B7A | N.D. | GCA_000725265.1 | *Escherichia coli* |  |
| BL21 (TaKaRa) | 93 | GCA_000833145.1 | *Escherichia coli* |  |
| BL21(DE3) | 93 | GCA_000009565.2 | *Escherichia coli* |  |
| BL21(DE3) | 93 | GCA_000022665.2 | *Escherichia coli* |  |
| BL21‑Gold(DE3)pLysS AG | 93 | GCA_000023665.1 | *Escherichia coli* |  |
| C227‑11 | 678 | GCA_000986765.1 | *Escherichia coli* |  |
| C2566 | 93 | GCA_001559615.1 | *Escherichia coli* |  |
| C3026 | 10 | GCA_001559675.1 | *Escherichia coli* |  |
| C3029 | 93 | GCA_001559635.1 | *Escherichia coli* |  |
| C321.deltaA | 10 | GCA_000474035.1 | *Escherichia coli* |  |
| C41(DE3) | 93 | GCA_000830035.1 | *Escherichia coli* |  |
| C43(DE3) | 93 | GCA_001039415.1 | *Escherichia coli* |  |
| CB9615 | 335 | GCA_000025165.1 | *Escherichia coli* |  |
| CD306 | 131 | GCA_001513615.1 | *Escherichia coli* |  |
| CE10 | 62 | GCA_000227625.1 | *Escherichia coli* |  |
| CFSAN029787 | 99 | GCA_001007915.1 | *Escherichia coli* |  |
| CFT073 | 73 | GCA_000007445.1 | *Escherichia coli* |  |
| CI5 | 5082 | GCA_000971615.1 | *Escherichia coli* |  |
| clone D i14 | 73 | GCA_000233895.1 | *Escherichia coli* |  |
| clone D i2 | 73 | GCA_000233875.1 | *Escherichia coli* |  |
| CQSW20 | 1060 | GCA_001455385.1 | *Escherichia coli* |  |
| DH1 #1 | 1060 | GCA_000023365.1 | *Escherichia coli* |  |
| DH1 #2 | 1060 | GCA_000270105.1 | *Escherichia coli* |  |
| DH1Ec095 | 1060 | GCA_001183645.1 | *Escherichia coli* |  |
| DH1Ec104 | 1060 | GCA_001183665.1 | *Escherichia coli* |  |
| DH1Ec169 | 1060 | GCA_001183685.1 | *Escherichia coli* |  |
| DHB4 | 10 | GCA_001559655.1 | *Escherichia coli* |  |
| E2348/69 | 15 | GCA_000026545.1 | *Escherichia coli* |  |
| E24377A | 1132 | GCA_000017745.1 | *Escherichia coli* |  |
| EC4115 | 11 | GCA_000021125.1 | *Escherichia coli* |  |
| EC958 | 131 | GCA_000285655.3 | *Escherichia coli* | **G821/H2129** |
| ECC‑1470 | 847 | GCA_000831565.1 | *Escherichia coli* |  |
| Eco889 | 131 | GCA_001663475.1 | *Escherichia coli* |  |
| Ecol_448 | 131 | GCA_001618365.1 | *Escherichia coli* |  |
| Ecol_732 | 131 | GCA_001617565.1 | *Escherichia coli* | **I1121** |
| Ecol_743 | 131 | GCA_001618325.1 | *Escherichia coli* |  |
| Ecol_745 | 131 | GCA_001618345.1 | *Escherichia coli* |  |
| ECONIH1 | 648 | GCA_000784925.1 | *Escherichia coli* |  |
| EDL933 | 11 | GCA_000732965.1 | *Escherichia coli* |  |
| ER1821R | 10 | GCA_001663075.1 | *Escherichia coli* |  |
| ER2796 | 10 | GCA_000800215.1 | *Escherichia coli* |  |
| ER3413 | 10 | GCA_000800765.1 | *Escherichia coli* |  |
| ER3435 | N.D. | GCA_000974885.1 | *Escherichia coli* |  |
| ER3440 | N.D. | GCA_000974465.1 | *Escherichia coli* |  |
| ER3445 | N.D. | GCA_000974535.1 | *Escherichia coli* |  |
| ER3446 | N.D. | GCA_000974825.1 | *Escherichia coli* |  |
| ER3454 | N.D. | GCA_000974405.1 | *Escherichia coli* |  |
| ER3466 | N.D. | GCA_000974575.1 | *Escherichia coli* |  |
| ER3475 | N.D. | GCA_000974865.1 | *Escherichia coli* |  |
| ER3476 | N.D. | GCA_000974505.1 | *Escherichia coli* |  |
| ETEC H10407 | 48 | GCA_000210475.1 | *Escherichia coli* |  |
| FRIK2069 | 11 | GCA_001651925.1 | *Escherichia coli* |  |
| FRIK2455 | 11 | GCA_001651965.1 | *Escherichia coli* |  |
| FRIK2533 | 11 | GCA_001651945.1 | *Escherichia coli* |  |
| G749 | 131 | GCA_001566635.1 | *Escherichia coli* |  |
| HS | 46 | GCA_000017765.1 | *Escherichia coli* |  |
| HUSEC2011 | 678 | GCA_000967155.1 | *Escherichia coli* |  |
| IAI1 | 1128 | GCA_000026265.1 | *Escherichia coli* | E650 |
| IAI39 | 62 | GCA_000026345.1 | *Escherichia coli* |  |
| IHE3034 | 95 | GCA_000025745.1 | *Escherichia coli* |  |
| JEONG‑1266 | 11 | GCA_001558995.2 | *Escherichia coli* |  |
| JJ1886 | 131 | GCA_000493755.1 | *Escherichia coli* |  |
| JJ1887 | 131 | GCA_001593565.1 | *Escherichia coli* |  |
| JJ1897 | 131 | GCA_001513655.1 | *Escherichia coli* |  |
| JJ2434 | 131 | GCA_001513635.1 | *Escherichia coli* |  |
| JW5437‑1 substr. MG1655 | 10 | GCA_001566335.1 | *Escherichia coli* |  |
| AG100 | 7415 | GCA_000981485.1 | *Escherichia coli* |  |
| BW25113 | 1996 | GCA_000750555.1 | *Escherichia coli* |  |
| BW2952 | 5967 | GCA_000022345.1 | *Escherichia coli* |  |
| DH10B | 4638 | GCA_000019425.1 | *Escherichia coli* |  |
| GM4792 #1 | 10 | GCA_001020945.2 | *Escherichia coli* |  |
| GM4792 #2 | 10 | GCA_001021005.2 | *Escherichia coli* |  |
| HMS174 | 10 | GCA_000953515.1 | *Escherichia coli* |  |
| MC4100 | 10 | GCA_000499485.1 | *Escherichia coli* |  |
| MDS42 | 1060 | GCA_000350185.1 | *Escherichia coli* |  |
| MG1655 #1 | 10 | GCA_000005845.2 | *Escherichia coli* |  |
| MG1655 #2 | 10 | GCA_000801205.1 | *Escherichia coli* |  |
| MG1655 #3 | 1060 | GCA_001308065.1 | *Escherichia coli* |  |
| MG1655 #4 | 10 | GCA_001544635.1 | *Escherichia coli* |  |
| MG1655_TMP32XR1 | 10 | GCA_001308125.1 | *Escherichia coli* |  |
| MG1655_TMP32XR2 | 10 | GCA_001308165.1 | *Escherichia coli* |  |
| RV308 | 10 | GCA_000952955.1 | *Escherichia coli* |  |
| W3110 | 10 | GCA_000010245.1 | *Escherichia coli* |  |
| KLY | 10 | GCA_000725305.1 | *Escherichia coli* |  |
| KO11 | 1079 | GCA_000147855.3 | *Escherichia coli* |  |
| KO11FL | 1079 | GCA_000258025.1 | *Escherichia coli* |  |
| LF82 | 135 | GCA_000284495.1 | *Escherichia coli* |  |
| LY180 | 1079 | GCA_000468515.1 | *Escherichia coli* |  |
| MNCRE44 | 131 | GCA_000931565.1 | *Escherichia coli* |  |
| MRE600 | N.D. | GCA_001542675.2 | *Escherichia coli* |  |
| MVAST0167 | 131 | GCA_001566655.1 | *Escherichia coli* |  |
| NA114 | 131 | GCA_000214765.2 | *Escherichia coli* |  |
| NCM3722 | 10 | GCA_001043215.1 | *Escherichia coli* |  |
| NGF1 | 998 | GCA_001660585.1 | *Escherichia coli* |  |
| Nissle 1917 | 73 | GCA_000714595.1 | *Escherichia coli* |  |
| NRG 857C | 135 | GCA_000183345.1 | *Escherichia coli* |  |
| P12b | 10 | GCA_000257275.1 | *Escherichia coli* |  |
| PCN033 | 5147 | GCA_000219515.3 | *Escherichia coli* | D2373 |
| PCN061 | 46 | GCA_001029125.1 | *Escherichia coli* |  |
| REL606 | 93 | GCA_000017985.1 | *Escherichia coli* |  |
| RM12579 | 335 | GCA_000245515.1 | *Escherichia coli* |  |
| RM12581 | 32 | GCA_000671295.1 | *Escherichia coli* |  |
| RM12761 | 6130 | GCA_000662395.1 | *Escherichia coli* |  |
| RM13514 | 32 | GCA_000520035.1 | *Escherichia coli* |  |
| RM13516 | 6130 | GCA_000520055.1 | *Escherichia coli* |  |
| RM9387 | 2773 | GCA_000801165.1 | *Escherichia coli* |  |
| RR1 | 10 | GCA_001276585.1 | *Escherichia coli* |  |
| RS218 | N.D. | GCA_000800845.2 | *Escherichia coli* |  |
| S51 | 7060 | GCA_001660565.1 | *Escherichia coli* |  |
| S88 | 95 | GCA_000026285.1 | *Escherichia coli* |  |
| Sakai | 11 | GCA_000008865.1 | *Escherichia coli* |  |
| Sanji | 167 | GCA_001610755.1 | *Escherichia coli* | **Z821** |
| Santai | 1011 | GCA_000827105.1 | *Escherichia coli* |  |
| SaT040 | 131 | GCA_001566615.1 | *Escherichia coli* |  |
| SE11 | 156 | GCA_000010385.1 | *Escherichia coli* |  |
| SE15 | 131 | GCA_000010485.1 | *Escherichia coli* |  |
| SEC470 | 48 | GCA_000987875.1 | *Escherichia coli* |  |
| SF‑088 | 95 | GCA_001280325.1 | *Escherichia coli* |  |
| SF‑166 | 95 | GCA_001280385.1 | *Escherichia coli* |  |
| SF‑173 | 95 | GCA_001280405.1 | *Escherichia coli* |  |
| SF‑468 | 95 | GCA_001280345.1 | *Escherichia coli* |  |
| SMS‑3‑5 | 354 | GCA_000019645.1 | *Escherichia coli* | E2372 |
| SQ110 | 10 | GCA_000988425.1 | *Escherichia coli* |  |
| SQ171 | 10 | GCA_000988445.1 | *Escherichia coli* |  |
| SQ2203 | 10 | GCA_000988465.1 | *Escherichia coli* |  |
| SQ37 | 10 | GCA_000988355.1 | *Escherichia coli* |  |
| SQ88 | 10 | GCA_000988385.1 | *Escherichia coli* |  |
| SRCC 1675 | 11 | GCA_001612495.1 | *Escherichia coli* |  |
| SS17 | 11 | GCA_000730345.1 | *Escherichia coli* |  |
| SS52 | 11 | GCA_000803705.1 | *Escherichia coli* |  |
| ST2747 | 6131 | GCA_000599665.1 | *Escherichia coli* |  |
| ST2747 | 6131 | GCA_000599685.1 | *Escherichia coli* |  |
| ST2747 | 6131 | GCA_000599705.1 | *Escherichia coli* |  |
| ST540 #1 | 540 | GCA_000597845.1 | *Escherichia coli* |  |
| ST540 #2 | 540 | GCA_000599625.1 | *Escherichia coli* |  |
| ST540 #3 | 540 | GCA_000599645.1 | *Escherichia coli* |  |
| ST648 | 648 | GCA_001485455.1 | *Escherichia coli* |  |
| TW14359 | 11 | GCA_000022225.1 | *Escherichia coli* |  |
| uk_P46212 | 131 | GCA_001469815.1 | *Escherichia coli* |  |
| UM146 | 643 | GCA_000148605.1 | *Escherichia coli* |  |
| UMNK88 | 100 | GCA_000212715.2 | *Escherichia coli* |  |
| UTI89 | 95 | GCA_000013265.1 | *Escherichia coli* |  |
| VR50 | 10 | GCA_000968515.1 | *Escherichia coli* |  |
| W #1 | 1079 | GCA_000184185.1 | *Escherichia coli* |  |
| W #2 | 1079 | GCA_000258145.1 | *Escherichia coli* |  |
| WS4202 | 11 | GCA_001307215.1 | *Escherichia coli* |  |
| Xuzhou21 | 11 | GCA_000262125.1 | *Escherichia coli* |  |
| YD786 | 410 | GCA_001442495.1 | *Escherichia coli* |  |
| ZH063 | 131 | GCA_001577325.1 | *Escherichia coli* |  |
| ZH193 | 131 | GCA_001566675.1 | *Escherichia coli* |  |
| 24 | 4633 | GCA_001514575.1 | *Escherichia albertii* |  |
| jun‑51 | 678 | GCA_001514595.1 | *Escherichia albertii* |  |
| 94389 | 11 | GCA_001514625.1 | *Escherichia albertii* |  |
| 20H38 | 6057 | GCA_001514555.1 | *Escherichia albertii* |  |
| CB10113 | 6054 | GCA_001514825.1 | *Escherichia albertii* |  |
| CB9791 | 6049 | GCA_001514845.1 | *Escherichia albertii* |  |
| E2675 | 2683 | GCA_001514865.1 | *Escherichia albertii* |  |
| EC03‑127 | 6052 | GCA_001514885.1 | *Escherichia albertii* |  |
| EC03‑195 | 4947 | GCA_001514905.1 | *Escherichia albertii* |  |
| EC05‑160 | 3762 | GCA_001514925.1 | *Escherichia albertii* |  |
| EC05‑44 | 6058 | GCA_001514945.1 | *Escherichia albertii* |  |
| EC05‑81 | 6055 | GCA_001514965.1 | *Escherichia albertii* |  |
| EC06‑170 | N.D. | GCA_001549955.1 | *Escherichia albertii* |  |
| HIPH08472 | N.D. | GCA_001514985.1 | *Escherichia albertii* |  |
| K7394 | 10 | GCA_001515005.1 | *Escherichia albertii* |  |
| K7744 | 10 | GCA_001515025.1 | *Escherichia albertii* |  |
| K7756 | 10 | GCA_001515045.1 | *Escherichia albertii* | A2361 |
| KF1 | 10 | GCA_000512125.1 | *Escherichia albertii* |  |
| KU20110014 | 3762 | GCA_001515065.1 | *Escherichia albertii* |  |
| LMG20976 | 383 | GCA_000759775.1 | *Escherichia albertii* |  |
| NIAH_Bird_13 | 3762 | GCA_001514645.1 | *Escherichia albertii* |  |
| NIAH_Bird_16 | 6056 | GCA_001514665.1 | *Escherichia albertii* |  |
| NIAH_Bird_2 | 4606 | GCA_001514685.1 | *Escherichia albertii* |  |
| NIAH_Bird_23 | 6059 | GCA_001514705.1 | *Escherichia albertii* |  |
| NIAH_Bird_24 | 4634 | GCA_001514725.1 | *Escherichia albertii* |  |
| NIAH_Bird_25 | 5967 | GCA_001514745.1 | *Escherichia albertii* |  |
| NIAH_Bird_26 | N.D. | GCA_001514765.1 | *Escherichia albertii* |  |
| NIAH_Bird_5 | 4736 | GCA_001514785.1 | *Escherichia albertii* |  |
| NIAH_Bird_8 | 2700 | GCA_001514805.1 | *Escherichia albertii* |  |
| TW07627 | 383 | GCA_000155105.1 | *Escherichia albertii* |  |
| TW08933 | 1763 | GCA_000208425.2 | *Escherichia albertii* |  |
| TW15818 | N.D. | GCA_000208505.2 | *Escherichia albertii* |  |

### Supplemental Table S1: Strains and NCBI accession numbers of strains used in the construction of the phylogenetic tree.

A total of 178 *E. coli* and 32 *E. albertii* genome sequences were used to construct the phylogenetic tree. The closest match to the colistin‑resistant strains in this study is indicated. For strains in bold we were able to link mutations in *basRS* to colistin resistance. Strain E3090 carried the mobile colistin resistance gene *mcr1.1*. When multiple GenBank assemblies had the same strain name, a numerical indicator was added. For some strains, MLST typing was not possible, this is denoted with N.D. (not determined).
