## Supplemental Table S2 for "Non-clonal emergence of colistin resistance associated with mutations in the BasRS two-component system in *Escherichia coli* bloodstream isolates"

| **Primer** | **Sequence (5’‑3’)** | **Reference** |
| --- | --- | --- |
| BW25113 *glmS* Up | ATA TTC AGT CAA TTA CAA ACA TTA | This study |
| BW25113 *glmS* Down | CGA TCT TCT ACA CCG TTC | This study |
| Tn7R | CAC AGC ATA ACT GGA CTG ATT TC | (S1) |
| Tn7R inward | GAA ATC AGT CCA GTT ATG CTG TG | This study |
| Tn7L | ATT AGC TTA CGA CGC TAC ACC C | (S1) |
| Tn7L inward | GGG TGT AGC GTC GTA AGC TAA T | This study |
| BasRS sequencing Fwd | AAA GCC CGT ATC CGC AC | This study |
| BasRS sequencing Rev | GAT CTC ACG CAT GAT GTG GC | This study |
| BasRS deletion BW27848 check Fwd | CAA ACG CAA CAC TAT TCA CAA GAC | This study |
| BasRS deletion BW27848 check Rev | ATC TCT GAC GCG CAT ACT CTC | This study |
| pGRG36 check with Tn7L | ATA TGC ACA GAT GAA AAC GGT G | This study |
| BasRS promoter Fwd | CTT CCT CTA CTG CAT CTG GG | This study |
| BasRS promoter E650 Fwd | CTT CCT CTA CTG CAT TTG GG | This study |
| BasRS promoter A2361 Fwd | TTT TCT CTA CTG CAT CTG GG | This study |
| BasRS promoter Rev | CAC GGT GTT TCC ATC GA | This study |
| BasRS promoter A2361 Rev | CAC GGT ATT TCC ATC AA | This study |
| BasRS genes Fwd | ATG AAA ATT CTG ATT GTT GAA GA | This study |
| BasRS genes Rev | GTT CAG CGT GCT GGT GGT | This study |
| BasRS genes A2361 Rev | ATT CAG CGT GCT GGT CGT | This study |
| BasRS genes overlap Fwd | TGC GCA CTT TGT TCG ATG GAA ACA CCG TGA TGA AAA TTC TGA TTG TTG AAG ACG AT | This study |
| BasRS genes overlap A2361 Fwd | TGC GCA CTT TGT TTG ATG GAA ATA CCG TGA TGA AAA TTC TGA TTG TTG AAG ACG AT | This study |
| BasRS promoter NotI Fwd | TAT CCT GCG GCC GCC TTC CTC TAC TGC ATC TGG G | This study |
| BasRS promoter NotI E650 Fwd | TAT CCT GCG GCC GCC TTC CTC TAC TGC ATT TGG G | This study |
| BasRS promoter NotI A2361 Fwd | TAT CCT GCG GCC GCT TTT CTC TAC TGC ATC TGG G | This study |
| BasRS genes XhoI Rev | TAT CCC CTC GAG GTT CAG CGT GCT GGT GGT | This study |
| BasRS genes XhoI A2361 Rev | TAT CCT CTC GAG ATT CAG CGT GCT GGT CGT | This study |
| BasRS promotor NotI short Fwd | TAT CCT GCG GCC GCC | This study |
| BasRS promotor NotI short A2361 Fwd | TAT CCT GCG GCC GCT | This study |
| BasRS promotor XhoI short Rev | TAT CCC CTC GAG GTT CA | This study |
| BasRS promotor XhoI short A2361 Rev | TAT CCT CTC GAG GTT CA | This study |
| I1121 BasS R10L mutagenesis Fwd | CGA CCA ATA TCG CTG CGC CAA CGG CTG | This study |
| I1121 BasS R10L mutagenesis Rev | GCG CAG AAA ACG CAT CAG ATT CAA TTA G | This study |
| H2129 BasR S53G mutagenesis Fwd | AGC CTG GTG GTA CTG GAT TTA GGC TTA CCC GAT G | This study |
| H2129 BasR S53G mutagenesis Rev | GTA ATG ACC GGC TTC AAG GCT TTG TTC CGC | This study |
| G821 BasS duplication mutagenesis Fwd | TTT CAT TAT CGA GCG TGC TGG | This study |
| G821 BasS duplication mutagenesis Rev | GGT TGT TTA CCG CTG ACG TC | This study |
| G821 BasS duplication check Fwd | ATC TGC TAT CAG GCG GTA CG | This study |
| G821 BasS duplication check Rev | GTT CGT CAT ACG AGG GGA GA | This study |
| Z821 BasS P159A mutagenesis Fwd | GAA CGC CAC TGG CGG GGG TGC GT | This study |
| Z821 BasS P159A mutagenesis Rev | GCA GTT CGT GCG CGA CGT CAG CGG TAA ACA A | This study |

### Supplemental Table S2: Sequences of primers used in this study.

Primers were used for all colistin‑resistant strains unless a specific strain name is provided in the description. In the primers used for inverse PCR site‑directed mutagenesis, the forward (fwd) primer carries the desired mutation, whilst the reverse (rev) primers complements the primer set for inverse PCR. The mutagenesis primers for strain G821 are designed to edit out the 162 nucleotide duplication in *basS*.
